# Interplay of septin amphipathic helices in sensing membrane-curvature and filament bundling

**DOI:** 10.1101/2020.05.12.090787

**Authors:** Benjamin L. Woods, Kevin S. Cannon, Amy S. Gladfelter

## Abstract

The curvature of the membrane defines cell shape. Septins are GTP-binding proteins that assemble into heteromeric complexes and polymerize into filaments at areas of micron-scale membrane curvature. An amphipathic helix (AH) domain within the septin complex is necessary and sufficient for septins to preferentially assemble onto micron-scale curvature. Here we report that the non-essential fungal septin, Shs1, also has an AH domain capable of recognizing membrane curvature. In mutants lacking a fully functional Cdc12 AH domain, the Shs1 AH domain becomes essential. Moreover, we find that the Cdc12 AH domain is also important for septin bundling, suggesting multiple functions for septin AH domains.

## Introduction

Cell shape is critical for function and can be thought of in terms of membrane curvature. Cell membranes range in curvatures from nanometer to micron scales. How cells use nanometer sized proteins to sense micron-scale membrane curvature is not well understood. One way cells solve this problem is through a class of cytoskeletal proteins called septins, which are conserved from yeast to humans (Field et al., 1996; Pan et al., 2007). Septins are GTP-binding proteins that form heteromeric complexes (Bertin et al., 2008; Low and Macara, 2006; Sirajuddin et al., 2007; Versele et al., 2004). These complexes associate with the plasma membrane, where they can polymerize into filaments and higher order assemblies such as bundles and rings (Bertin et al., 2008; DeMay et al., 2011; DeMay et al., 2009; John et al., 2007; Rodal et al., 2005; Sirajuddin et al., 2007). Higher order septin assemblies can act as scaffolds for signaling proteins, organize membrane properties and influence the organization of other cytoskeletal elements (Bridges et al., 2016; Clay et al., 2014; Gilden et al., 2012; Gladfelter et al., 2001; Lew and Reed, 1995; Longtine et al., 2000; Spiliotis, 2010). In many of these functions, higher order septin assemblies are built in the context of micron-scale curvature.

Septins can distinguish positive membrane curvatures at the micron-scale without any other cellular factors (Bridges et al., 2016). When purified septins are mixed with membranes at a range of different curvatures, septins polymerize into aligned filaments wrapped at an optimal curvature (Beber et al., 2019; Cannon et al., 2019; Tanaka-Takiguchi et al., 2009). However, individual septin complexes can distinguish membranes with micron-scale curvature which manifests through septin-membrane binding kinetics (Cannon et al., 2019). Septins, like other curvature sensitive proteins, possess an amphipathic helix (AH) domain (Cannon et al., 2019; Drin and Antonny, 2010; Kim et al., 2017). AH domains function by binding to lipid packing defects within curved membranes. In the budding yeast *Saccharomyces cerevisiae*, the amphipathic helix at the C-terminus of Cdc12 is necessary and sufficient for septins to distinguish between different curvatures of the plasma membrane. Thus, the ability of septins to sense membrane curvature occurs at multiple scales.

Here we report that Shs1, a non-essential mitotic septin, also has an AH domain within its C-terminal extension (CTE). Shs1 occupies the terminal position at either end of a subset of heteromers, adjacent to Cdc12, the only essential septin subunit to possess an AH domain in yeast (Cannon et al., 2019; Finnigan et al., 2015; Garcia et al., 2011). Like the Cdc12 AH domain, the Shs1 AH domain recognizes membrane curvature. In the absence of the Cdc12 AH domain, the Shs1 AH domain restores the ability of septins to distinguish between different curvatures. Moreover, Shs1 and its CTE are indispensable in mutants harboring an incomplete Cdc12 AH domain in cells. This suggests septin complexes can contain different numbers of curvature sensitive AH domains, which may allow for another layer of control of higher order septin assemblies.

## Results and Discussion

### A predicted AH domain of Shs1 can distinguish between different membrane curvatures *in vitro*

We reported previously that septin complexes harboring Cdc12 mutant protein, cdc12-6, where the AH domain is truncated, no longer could distinguish between different membrane curvatures (Cannon et al., 2019). In *cdc12-6* mutants, septins quickly disassemble from the bud neck at restrictive temperature. However, at permissive temperature septins remain localized to bud neck, which is a site of positive micron-scale membrane curvature in cells (Gladfelter et al., 2005). We reasoned that another, unidentified AH domain might enable septins to localize in these mutants at permissive temperature. Screening through protein sequences of all mitotically expressed septins in ascomycetes including *S. cerevisiae*, *A. gossypii*, and *K. lactis* revealed an AH domain within the CTE of Shs1 (Figure 1, B and C). Interestingly, the ascomycetes *N. crassa*, *S. pombe*, and *A. nidulans* lack an Shs1 homolog all together, whereas *C. albicans* Shs1 does not have an AH. Despite sharing relatively low homology (∼65%) among AH-containing Shs1 polypeptides (Figure 1, A), AH domains are nearly identical (Figure 1 B-D). The physicochemical properties of AH domains have been shown to affect membrane binding and curvature sensing (Drin and Antonny, 2010; Drin et al., 2007). Interestingly, the charge and hydrophobic moment of the Shs1 AH are different than the Cdc12 AH (Figure 1, A and B). Together, these data suggest that the Shs1 AH domain might impart a distinct curvature preference when combined with Cdc12 in the same septin complex.

**Figure 1:**
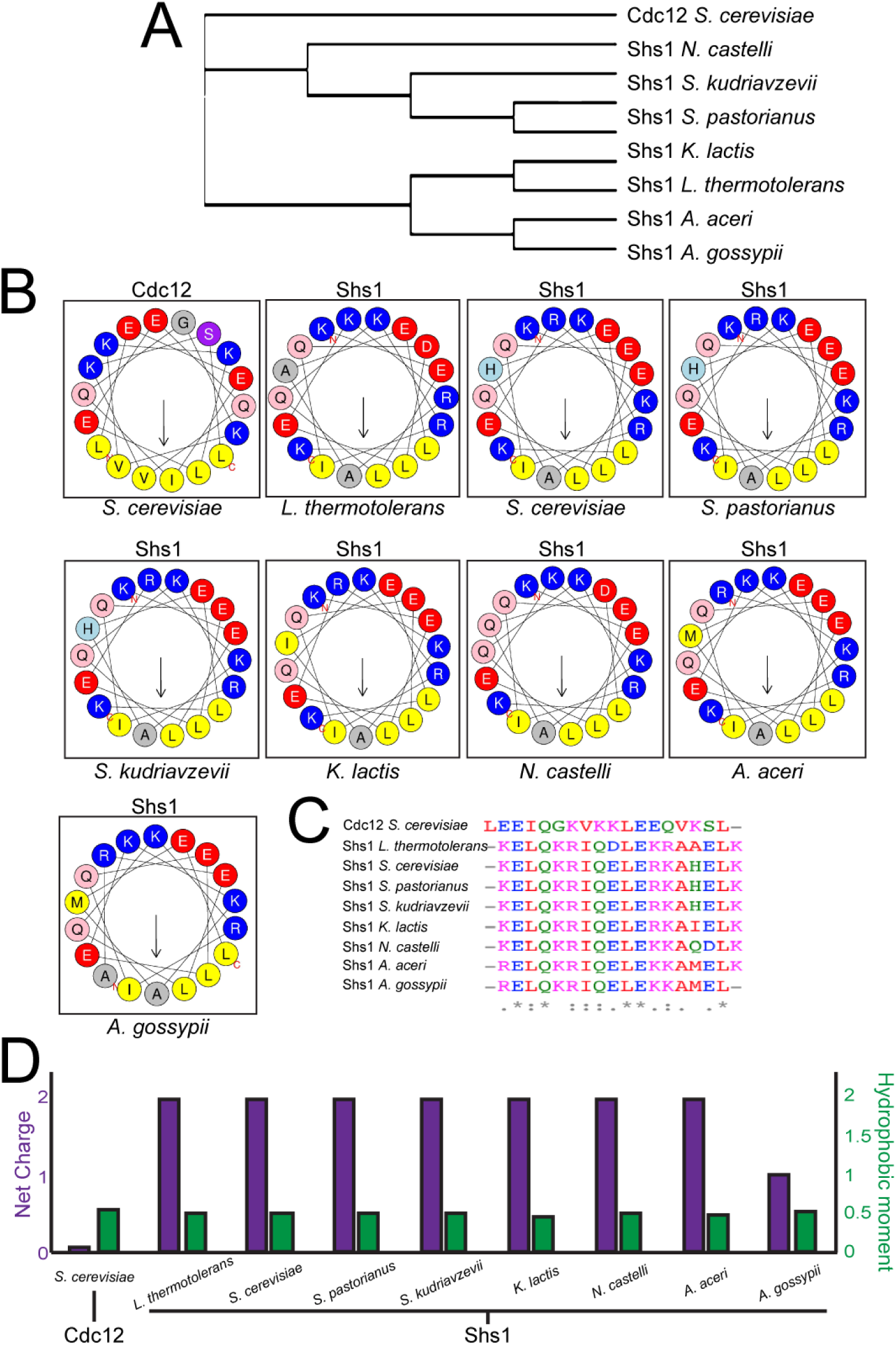
Shs1 contains a highly conserved amphipathic helix within its C-terminal domain. A. Cladogram constructed from multiple alignments of Shs1 primary sequences from various ascomycetes. B. Helical wheels representing amphipathic helices present in septin polypeptides screened ascomycetes. C. Sequence alignment of amphipathic helices. D. Net charge and hydrophobic moment of septin amphipathic helices.

To assess whether the *S. cerevisiae* Shs1 AH domain can distinguish between membrane curvatures, we purified a polypeptide with 2 copies of the AH of Shs1 to mimic the stoichiometry of the septin complex and mixed it with membrane-coated, silica beads ranging in diameter from 300 nm to 5 μm (curvatures of κ = 6.67 μm^−1^ to κ = 0.4 μm^−1^ respectively). The Shs1 AH domain adsorbed best to higher curvatures (although the preference between curvatures of 6.67 μm^−1^ and 2 μm^−1^ were not statistically significant), with almost no binding at lower curvatures (Figure 2A). Thus, the Shs1 AH can distinguish different curvatures and - despite the different chemical properties - has a similar membrane curvature preference to the Cdc12 AH (Cannon et al., 2019).

**Figure 2:**
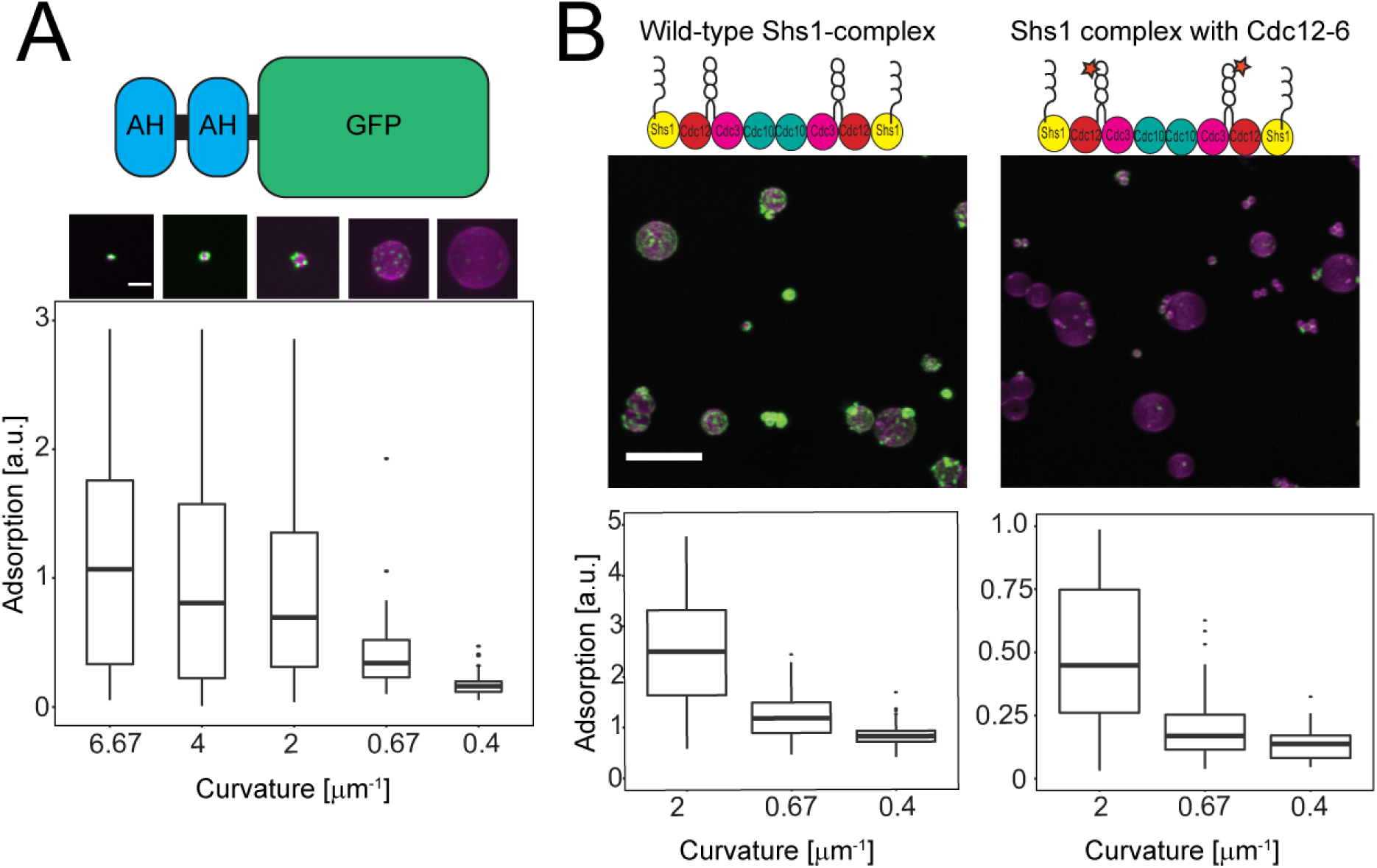
Shs1’s amphipathic helix is capable of binding micron-scale membrane curvatures. A. (Top) Maximum projection images of 2 µM 2x-Shs1AH-GFP (green) adsorbed onto curved supported lipid bilayers (magenta). Scale bar, 2 µm. Images contrasted identically. (Bottom) Box and whisker plot of 2x-Shs1AH-GFP adsorption onto different membrane curvatures. Black bars represent the median. Error bars are the standard deviation for N >10 beads at each curvature. B. (Top) Maximum projection images of 100 nM wild-type Shs1 complex (green, panel 1) or Shs1-Cdc12-6 complex (green, panel 2) on curved supported lipid bilayers (magenta). Scale bar, 10 µm. Images are contrasted identically. (Bottom) Box and whisker plot quantifying adsorption onto different curvatures. Black bars represent the median. Error bars are the standard deviation for N >70 beads at each curvature.

Why does Shs1 harbor a curvature-sensing AH domain? Within the septin complex Cdc12 resides at the penultimate position, with the terminal subunit being either Cdc11 or Shs1 (Bertin et al., 2008; Versele et al., 2004) and there is no evidence of “chimeric” octamers with Cdc11/Shs1 on either end (Garcia et al., 2011; Khan et al., 2018; Weems and McMurray, 2017). Thus, Cdc11-capped complexes possess two AH domains, spaced apart by ∼24 nm (the distance between the Cdc12 subunits) while Shs1-capped complexes have four AH domains, altering the valency and spacing of AHs within assemblies. This raises the possibility that septin curvature preference could be different when both AHs are combined in the same complex.

To test whether the Shs1 AH domain alters the curvature preference of septins, we purified Shs1-capped octamers and mixed them with different-sized membrane coated beads. Shs1-capped octamers displayed a similar membrane curvature preference to that of Cdc11-capped octamers with the highest adsorbance on beads 1 µm in diameter (κ = 2 μm^−1^) (Figure 2B) (Cannon et al., 2019). This is consistent with the behavior of tandem AH domains (Figure 2A and Cannon et al., 2019) and our previous observation where Cdc11-and Shs1-capped octamers were mixed and preferentially adsorbed onto membrane curvatures 1 µm in diameter (Khan et al., 2018). It was unclear why addition of two more AH domains did not alter membrane-curvature sensitivity. One possibility is that the second pair of AHs in the complex help stabilize membrane interactions, while not changing the curvature preference. This makes sense as Shs1 complexes have diminished ability to form filaments to provide avidity in membrane association.

Next, we tested if Shs1-capped complexes could restore membrane curvature sensitivity of cdc12-6 heteromeric-complexes in which the AH domain is truncated. Consistent with the Shs1 AH having a similar curvature preference as the Cdc12 AH, septin complexes harboring cdc12-6 and capped with Shs1 preferentially adsorbed onto similar membrane curvatures as complexes harboring wild-type Cdc12 (Figure 2B). Collectively, these data suggest that within a septin complex, the AH domains of Shs1 and Cdc12 sense membrane curvature at the same scale.

Despite restoration of curvature sensing, the adsorption of cdc12-6 Shs1-capped octamers were substantially lower than Cdc12-wt Shs1-capped octamers. This contrasts with cdc12-6 Cdc11-capped octamers which lack curvature sensitivity, adsorbing nonspecifically to all curvatures tested (Cannon et al., 2019). The differential adsorption of cdc12-6 octamers with either Cdc11 or Shs1 has several implications. First, it suggests the enhanced, non-specific binding of cdc12-6 complexes is dependent on Cdc11, suggesting a strong membrane-interaction surface in Cdc11. Second, given Shs1’s limited ability to polymerize, cdc12-6 Shs1-capped complexes may be less stably associated with the membrane and with an even greater handicap to form filaments, further reducing overall adsorption. Nonetheless, these data suggest that the Shs1 AH is a functional curvature sensor.

### The Shs1 CTE and AH domain are required for normal septin function in the *cdc12-6* mutant

As Shs1 harbors an AH domain, it is possible that Shs1 becomes critical for septin localization at the bud neck in *cdc12-6* strains at permissive temperature. If the ability of septins to differentiate membrane curvatures is essential, and either the Cdc12 or Shs1 AH domain can mediate curvature sensing, then compromising both would be synthetic lethal. Consistent with this hypothesis, it had been previously reported that deletion of *SHS1* in a *cdc12-6* background is inviable (Finnigan et al., 2015), which was corroborated by tetrad dissection in our strain background at 24°C (Figure 3B). To distinguish whether it was the Shs1 AH domain that was essential in *cdc12-6* mutants, we generated a panel of C-terminal Shs1 truncations in the *cdc12-6* background (Figure 3A). C-terminal truncations of Shs1 that begin immediately *after* the AH domain (*shs1^Δ506-551^* and *shs1^Δ508-551^*) were viable in the *cdc12-6* background at 24°C (Figure 3B). However, some double mutants had classic septin mutant phenotypes with elongated buds and often failed to separate from the mother (Fig 2C). This may be due to the proximity of the truncation to the AH domain. *cdc12-6* mutants expressing Shs1 truncation beginning 15 residues *after* the AH domain were fully viable without septin defects (Figure 3, B and C). *cdc12-6* mutants expressing a more extensive Shs1 truncation (*shs1^Δ488-551^*) or an AH deletion (*shs1^ΔAH^*) were sick, displaying fully penetrant septin defects at 24°C (Figure 3, B and C). Truncating the entire Shs1 CTE (*shs1^Δ341-551^*) was synthetic lethal with *cdc12-6* (Figure 3B). These data indicate that the AH domain within the CTE is necessary for proper septin function in *cdc12-6* mutants but that the full CTE is required for viability.

**Figure 3.**
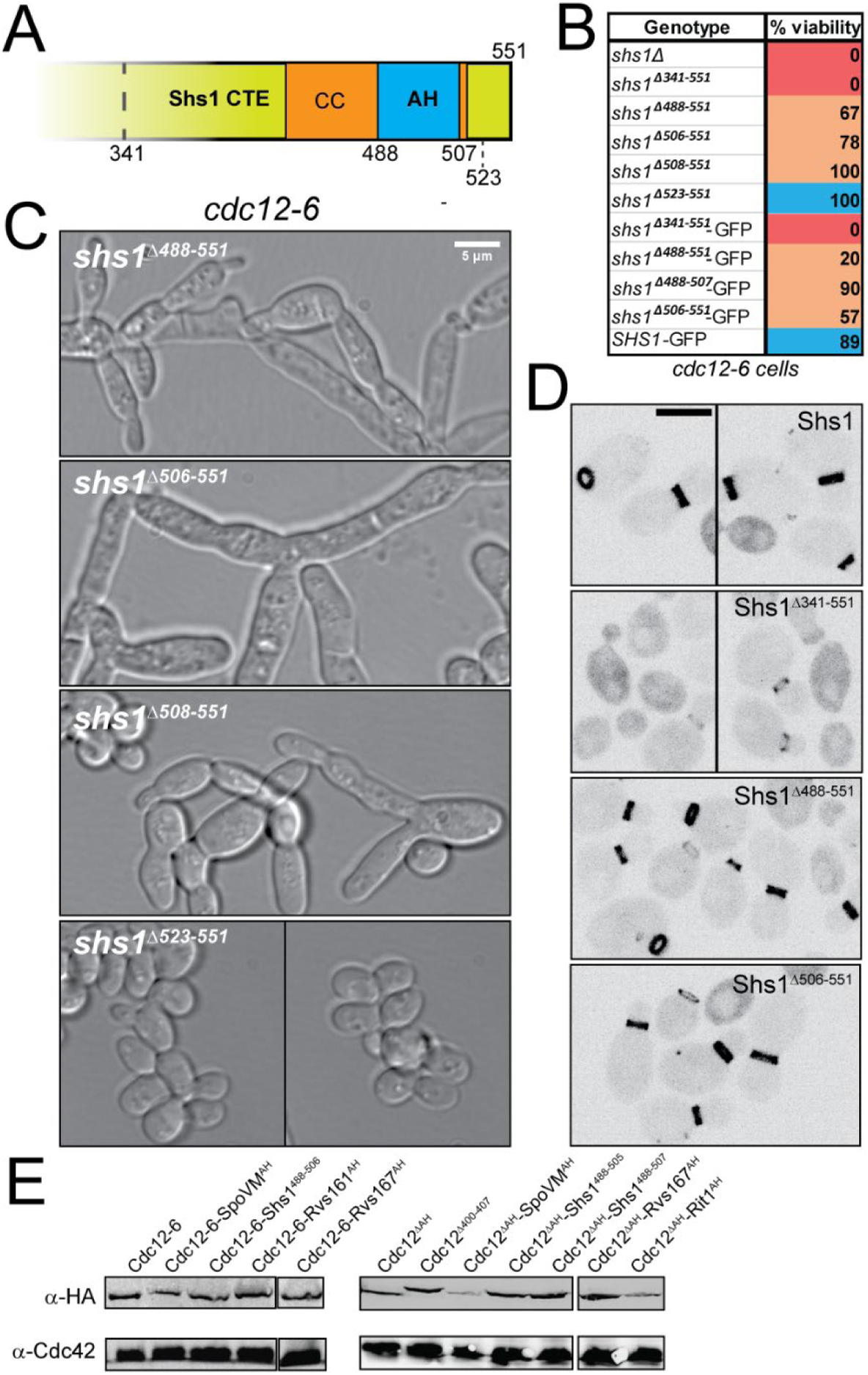
Genetic analyses of Shs1 and Cdc12 AH domains. A. *S. cerevisiae* Shs1 CTE with a predicted AH domain within the coiled coil. Truncation sites within Shs1 CTE demarcated. B. Viability of *cdc12-6* cells expressing indicated *SHS1* alleles expressed from the endogenous locus based on tetrad dissections at permissive temperature (24°C). Genotypes labeled in blue appeared normal, without obvious septin defects. Genotypes in orange were sick, with partially or fully penetrant septin defects. Genotypes in red were inviable. C. DIC images of *cdc12-6 shs1* mutants (see B) at 24°C. Scale bar, 5 μm. D. Heterozygous diploids expressing the indicated Shs1 protein fused to GFP from the *SHS1* locus. Scale bar, 5 μm. E. Western blot comparing expression of indicated Cdc12 chimeras fused to 3xHA epitope in heterozygous diploids.

It was possible that truncating Shs1 reduced Shs1 incorporation into septin complexes. To rule this out, we generated GFP tagged Shs1 truncations expressed from the endogenous *SHS1* locus to assess Shs1 expression and localization. In wild-type *CDC12* cells Shs1 truncations (with the exception of Shs1^Δ341-551^-GFP) were expressed at similar levels to wild-type Shs1-GFP and localized to the bud neck normally (Figure 3D). This suggests these truncations do not interfere with Shs1 incorporation. Shs1^Δ341-551^-GFP, in which the entire CTE is truncated, was expressed normally in wild-type cells, but localized predominantly in the cytoplasm with reduced signal at the bud neck (Figure 3D). The increased cytoplasmic distribution of Shs1^Δ341-551^-GFP provides an explanation for the observed synthetic lethality with *cdc12-6* mutants. In sum these data suggest that removal of the AH does not block Shs1 incorporation into septin complexes.

We reasoned that if the main function of the Cdc12 AH domain is to sense membrane curvature, then swapping it with other AH domains might be sufficient to restore cdc12-6 function. We generated chimeric strains where different AH domains were fused to the C-terminus of either cdc12-6 or Cdc12 lacking its entire AH domain (Cdc12^ΔAH^). Chimeric Cdc12 included AH domains from Shs1, Rvs161, Rvs167, *S. pombe* RitC, and *B. subtilus* SpoVM (Bendezu et al., 2015; Gill et al., 2015; Youn et al., 2010). Surprisingly, none of these chimeric strains were viable even at permissive temperature as assessed by tetrad dissection. This is despite that Cdc12 AH domain chimeras were expressed normally relative to wild-type Cdc12 (Figure 3E). Moreover, we could discount the possibility that the 3xHA tag adjacent to the AH domain was responsible for the lethality since the 3xHA tag adjacent to the Cdc12 AH domain (Cdc12^Δ400-407^) had no effect on viability (Figure 3B). These data indicate that the Cdc12 AH domain cannot be simply swapped for another AH domain, even if chimeric AH domain recognizes similar curvatures (Figure 2A) (Gill et al., 2015).

Why are *cdc12^ΔAH^* mutants inviable even in the presence of wild-type *SHS1*? It is possible that cdc12-6 containing octamers *in vivo* maintain some minimal ability to localize to membrane curvature that can be supplemented by Shs1. Although this contrasts with the *in vitro* data (Cannon et al., 2019). Another possibility is that cdc12^ΔAH^ does not fold properly or incorporate into octamers efficiently, rendering it non-functional. However, cdc12^ΔAH^ was expressed at wild-type levels (Figure 2E) suggesting it is not degraded and that is incorporated into septin complexes. A third possibility is that Shs1 does not efficiently cap cdc12^ΔAH^ septin oligomers.

### The Cdc12 AH domain inhibits septin bundling

Why are Cdc12-AH domain chimeras non-functional? And why are *cdc12-6* mutants sensitive to Shs1 CTE truncations, including ones that maintain the Shs1 AH? We reasoned there might be an additional, essential function to the Cdc12 AH domain or a gain-of-function that occurs when the curvature sensitivity of the AH domain is lost. A clue comes from yeast cells exposed to pheromone, where in *cdc12-6* mutants septins no longer localize into filaments in the shmoo, and typically remain cytoplasmic (Giot and Konopka, 1997; Longtine et al., 1998). In a subset of cells, we observed aberrant septin structures that appeared to be bundled “needles” and not in association with the cortex (Figure 4A). Needles were not observed in cycling, heterozygous diploids (*cdc12-6/CDC12*), possibly indicating that ectopic bundled needles are a recessive phenotype of *cdc12-6* mutants.

**Figure 4.**
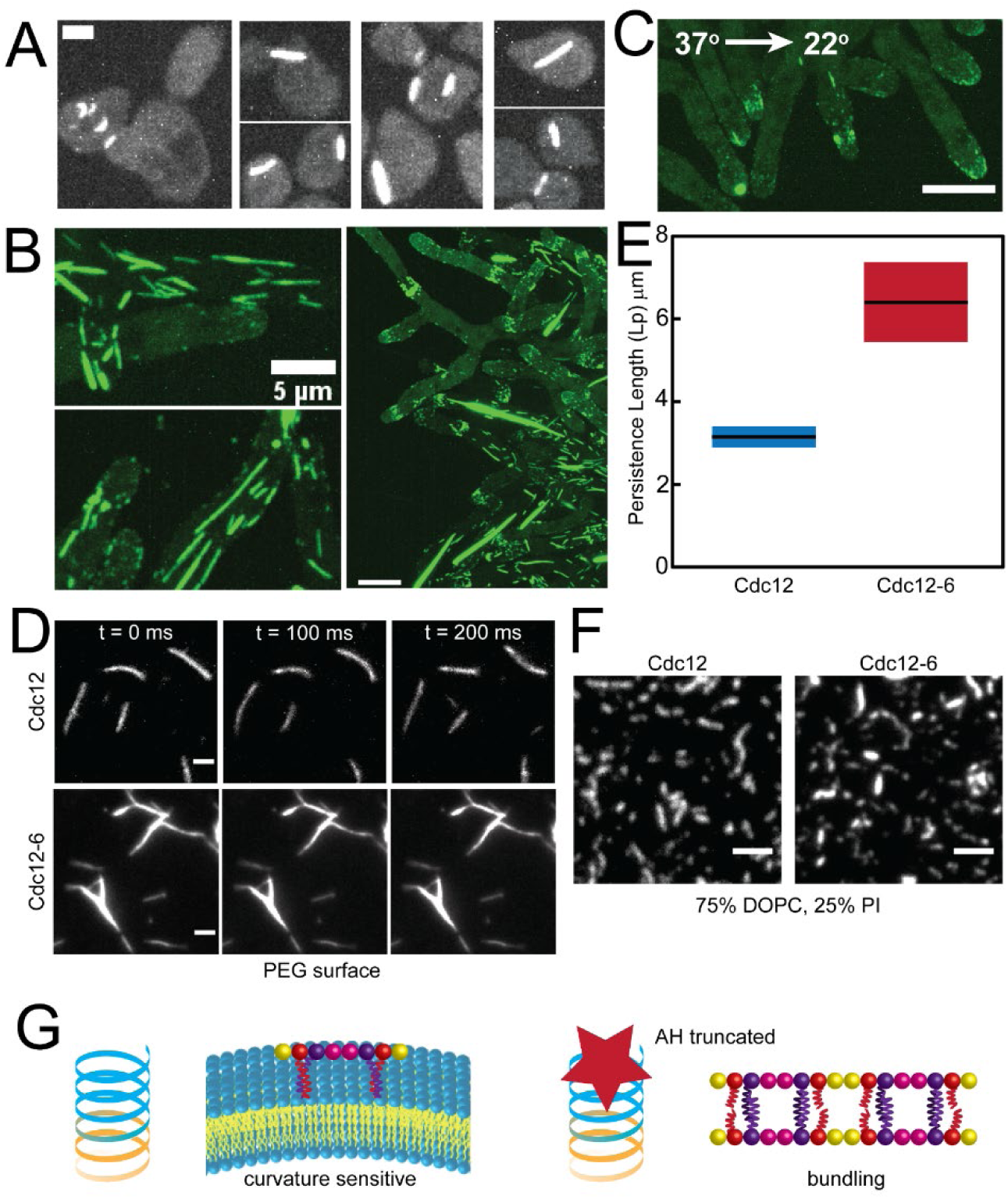
The Cdc12 AH domain inhibits septin bundling. A. Maximum projections (contrasted identically) of fixed cells with bundled “needles” in *cdc12-6 S. cerevisiae* mutants expressing Cdc3-mCherry treated with pheromone. Scale bar, 2 µm. B. Maximum projections of bundled “needles” in *A. gossypii* expressing Cdc12-6-GFP at 22°C. Scale bars, 5 μm (left panels) and 10 μm (right). C. Maximum projection of a *cdc12-6 A. gossypii* mutant at 22°C immediately after being shifted down from 37°C for one hour. Scale bar, 10 μm. D. TIRF images of wild-type (top) and cdc12-6 (bottom) septin filaments formed in solution. Scale bar, 10 µm. E. Persistence length as determined by the cosine correlation of wild-type and Cdc12-6 filament tangent angles in solution. Lines represent mean, bars represent range. F. TIRF images of 3 nM wild-type and cdc12-6 septins polymerized into filaments on planar supported lipid bilayers. Scale bar, 2 µm. G. The Cdc12 AH domain enables septins to discriminate micron-scale membrane curvature. Blocking the AH domain promotes septin bundling.

In *Ashbya cdc12-6* mutants, bundled needle-like structures were detectable in the cytoplasm of vegetatively growing cells, with some needles extending the entire length of hyphae (Figure 4B). Interestingly, *cdc12-6* septin needles are temperature sensitive and quickly disassemble when *Ashbya* mutants are shifted to 37°C (Figure 4C). Curiously, after shifting down from 37°C to 22°C, septins did not immediately reform into needles, but instead localized to hyphal tips (Figure 4C). This is consistent with a previous observation in budding yeast *cdc12-6* mutants in which after shifting down to permissive temperature, septins did not localize to the bud neck, but instead localized to the bud where polarity factors are localized (Gladfelter et al., 2005).

The existence of naturally occurring bundled septin structures in cells suggest that cdc12-6 bundles are not necessarily only a gain-of-function mutation but that the C-terminus of Cdc12 could be relevant for bundling septins under certain contexts (DeMay et al., 2009; Liu et al., 2019). However, given their abundance in *cdc12-6* mutants, the needles could act like a sponge, sequestering septin complexes from the cytoplasm into “function-less” structures. Consistent with this, recombinant cdc12-6 septin complexes (capped with Cdc11) form needles in solution that were less flexible than wild-type filaments (Figure 4, D and E, **Movie S1**). This indicates formation of needle-like structures is an intrinsic feature of cdc12-6 septin complexes and is not dependent on other cellular factors. Interestingly, when added on supported planar lipid bilayers, cdc12-6 octamers did not form bundled needles (Figure 4F), suggesting that membranes may play an inhibitory role in bundling. This is supported by the observation that needles in cells appeared to be within the cytosol away from the cortex.

The needle-like septin structures in *cdc12-6* mutants suggest the AH domain of Cdc12 may also have a role in septin bundling (Figure 4G). Truncation of the Cdc12 AH domain is predicted to misfold the coiled-coil element at the C-terminus of Cdc12 (*MultiCoil*) (Wolf et al., 1997). Within the septin complex, the coiled-coil element of Cdc12 is thought to associate with the coiled-coil element of Cdc3 in parallel (Bertin et al., 2008; Versele et al., 2004). Homologous coiled-coil elements can dimerize or even oligomerize into larger coiled-coil structures in the absence of the preferred binding partner (Harbury et al., 1993; Lupas and Bassler, 2017). We speculate that free Cdc3 coiled-coil elements could oligomerize in the absence of the Cdc12 coiled-coil elements to promote bundling. If coiled-coil misfolding is exacerbated in mutants lacking the AH, it could be another explanation as to why *cdc12^ΔAH^* mutants are inviable. In *Ashbya*, dispersed septin filaments near the tips of hyphae coalesce into discrete bundled structures along the hyphal body (DeMay et al., 2009). This transition is dependent on the kinase Gin4, which is predicted to interact with the coiled-coil element of Cdc3 and phosphorylates Shs1 in *S. cerevisiae* (Longtine et al., 1998; Mortensen et al., 2002). The Cdc3/Cdc12 coiled-coil may act like togglable switch, whose disassembly - either through phosphorylation or Cdc12 AH domain sequestration - could promote septin bundling through Cdc3 coiled-coil oligomerization.

### Conclusion

Here we report that an additional AH domain in the non-essential septin, Shs1, can recognize membrane-curvature. Shs1 and its CTE harboring the AH domain become essential in mutants lacking a fully functional AH domain in Cdc12 (*cdc12-6*). cdc12-6 induces filament bundling into non-physiological needle-like structures, possibly exacerbating its phenotype. This may indicate a role for the AH domain in septin bundling. Future research should investigate how the Shs1 AH domain might influence septin-membrane binding affinity and diffusion in the membrane. The role of AH domains in septin biochemistry and biology is only beginning to be understood and is an exciting area of future study.

## Materials and Methods

### Yeast strain construction and culturing

Standard molecular genetic techniques were used to generate and culture yeast strains in this study. All yeast strains used in this study are listed in **Table S1**. The temperature sensitive *cdc12-6* mutant was used assess the function of Shs1 truncations. We received a *bar1Δ* strain and a diploid strain heterozygous for *shs1Δ* and *cdc12-6* as gifts (D. Lew, Duke University). We generated the untagged Shs1 truncations (Shs1^Δ341-551^, Shs1^Δ488-551^, Shs1^Δ488-507^, Shs1^Δ506-551^, Shs1^Δ508-551^, and Shs1^Δ523-551^) and GFP-tagged Shs1 constructs (Shs1-GFP, Shs1^Δ341-551^-GFP, Shs1^Δ488-551^-GFP, Shs1^Δ506-551^-GFP) using the PCR-based C-terminal modification method (Longtine et al., 1998). We generated Cdc12-6, Cdc12^ΔAH^ AH-chimeric (SpoVM^AH^, Shs1^488-505^, Shs1^488-507^, Rvs161^AH^, Rvs167^AH^, *S. pombe* Rit1^AH^) and Cdc12^Δ400-407^ constructs with an HA-tag using the PCR-based C-terminal modification method mentioned above. PCR vectors were integrated at their endogenous loci (*SHS1* or *CDC12*, respectively) to generate heterozygous diploids. All transformants were verified for targeted integration by PCR amplification of genomic DNA.

Cells were grown to mid-log phase at 24°C in Complete Synthetic Medium (CSM, Sunrise Science Products) supplemented with 0.67% yeast nitrogen base, 2% dextrose, and 0.01% adenine, then harvested. Cells were mounted onto CSM + 2% dextrose agarose (2%) pads prior to imaging. For experiments using yeast pheromone, mid-log phase cultures of *bar1Δ* cells were treated with 50 nM α-factor (GenWay Biotech) for two hours at 24°C before fixation with 3.7% formaldehyde for 10 minutes at room temperature before imaging.

### *Ashbya gossypii* culturing

The *A. gossypii* strain harboring the *cdc12-6-*GFP allele at the endogenous *CDC12* locus (AG884; *AgCDC12:cdc12-6-*GFP:*GEN leu2Δ thr4Δ*) was described before (Cannon et al., 2019). *A gossypii* was grown from spores in full medium for 16 hours at 24°C before harvesting mycelial cells unless otherwise indicated. Cells were mounted on Low Fluorescence medium agarose (2%) pads for imaging.

### Cell microscopy

Confocal images were acquired using a spinning disc (Yokogawa W1) confocal microscope (Nikon Ti-82 stage) using a 100x Plan Apo 1.49 NA oil lens and a Prime 95B CMOS camera (Photometrics). Differential interference contrast (DIC) images were acquired on a custom-built microscope with an inverted Olympus IX-83 body equipped with an Olympus 60x 1.3 NA lens and an EMCCD camera (Andor iXon-888) (McQuilken et al., 2017). Live cell microscopy images were prepared using ImageJ (FIJI).

### Immunoblotting

For Western blot analysis, ∼2×10^7^ cells were harvested from mid-log phase cultures grown at 24°C and total protein was extracted by TCA precipitation as previously published (Keaton et al., 2008). Electrophoresis and Western blotting were performed as previously described (Bose et al., 2001). A polyclonal rabbit α-HA epitope antibody preparation was used at 1:2000 dilution (Rockland Immunochemicals). A monoclonal mouse α-Cdc42 antibody preparation (gift from P. Brennwald, UNC) was used at 1:500 dilution (Wu et al., 2015). Polyclonal goat antibodies against rabbit (Goat anti-Rabbit IgG DyLight 680 conjugated, Rockland Immunochemicals) and mouse (Goat anti-Mouse IgG DyLight 800 conjugated, Rockland Immunochemicals) were used at 1:10,000 dilution as secondary antibodies. Blots were imaged with an ODYSSEY infrared laser scanner (LI-COR Biosciences).

### Protein purification

Recombinant *S. cerevisiae* septin protein complexes were expressed from a duet expression system in BL21 (DE3) *E. coli* cells and purified as previously described (**Table S2**) (Bridges et al., 2016; Cannon et al., 2019). The Shs1 amphipathic helix conjugated to GFP was purified with a similar protocol as septins, however induced cultures were grown for four hours at 37°C instead of 24 hours at 22°C.

### Amphipathic helix constructs

The 2x AH(Shs1)-GFP construct has the primary sequence: <u>KELQKRIQELERKAHELK-</u>GSGSRSGSGS-<u>KELQKRIQELERKAHELK</u>-GSGSSR-GFP tag. The underline sequence represents the AH domain from Shs1. The glycine-serine repeats represent linker sequences between the AH domains and the GFP tag.

### Lipid mix preparation

Lipids (from Avanti Polar Lipids) were mixed in chloroform solvent at a ratio of 75 mole percent dioleoylphosphatidylcholine (DOPC) and 25 mole percent PI (liver, bovine) sodium salt in a glass cuvette. For lipid mix preparations to incubated on silica microspheres, trace amounts (>0.1%) of phosphatidylethanolamine-N-(lissamine rhodamine B sulfonyl) (RH-PE) were added. Lipid mixtures were then dried with argon gas to create a lipid film and stored under negative vacuum pressure overnight to evaporate trace chloroform. Lipids were rehydrated in Supported Lipid Bilayer buffer (300 mM KCl, 20 mM Hepes pH 7.4, 1 mM MgCl2) at 37°C to a final lipid concentration of 5 mM. Lipids were then resuspended over the course of 30 minutes at 37°C, with vortexing for 10 seconds every five minutes. Fully resuspended lipids were then bath sonicated at 2-minute intervals until solution clarified to yield small unilamellar vesicles (SUVs).

### Preparation of supported lipid bilayers

Lipid bilayers supported on silica microspheres were prepared as previously described (Bridges et al., 2016; Cannon et al., 2019). SUVs were adsorbed onto silica microspheres (Bangs Laboratories) by incubating 50 nM of lipid with different bead sizes (with a summed surface area of 440 mM^2^) for hour at room temperature with gentle rotation. Microspheres were pelleted, then resuspended in in Pre-Reaction buffer (33 mM KCl, 50 mM HEPES pH 7.4) to wash away excess SUVs. This wash step was repeated four additional times.

Planar supported lipid bilayers were prepared in a similar process as reported previously (Bridges et al., 2014). No. 1.5 coverslips were first cleaned with oxygen plasma (PE25-JW, Plasma Etch). Chambers for lipid bilayers cut from PCR tubes were glued (Norland optical adhesive, Thor Labs) onto cleaned coverslips. SUVs were incubated at 37°C in chambers on coverslips suspended in Supported Lipid Bilayer buffer with additional 1 mM CaCl_2_ for 20 minutes (1 mM final lipid concentration in 50 μl solution). After vesicle fusion, lipid bilayers were washed six times with 150 μl of Supported Lipid Bilayer buffer to rinse away excess SUVs. Immediately prior to adding septins, bilayers were further washed with a 150 μl of buffer containing 50 mM Hepes pH7.4, 1 mM BME, and 0.01% BSA six times.

### Measuring protein adsorption on lipid bilayers supported on silica microspheres

Experiments measuring the adsorption of septins and AH domains on different curvatures were performed as previously published (Cannon et al., 2019). Reactions (with a final buffer composition of 100 mM KCl, 50 mM Hepes pH 7.4, 1 mM BME, 0.1% methyl-cellulose, and 0.01% BSA) were incubated in a plastic chamber (Bridges and Gladfelter, 2016) glued onto a polyethylene glycol (PEG)-coated passivated coverslip for one hour to reach equilibria. Confocal images of fluorescent-tagged protein adsorbed onto curved supported bilayers on microspheres were acquired using a spinning disc (Yokogawa W1) confocal microscope (Nikon Ti-82 stage) using a 100x Plan Apo 1.49 NA oil lens and a Prime 95B CMOS camera (Photometrics). Raw images were analyzed using Imaris 8.1.2 software (Bitplane AG) as previously described (Cannon et al., 2019). Boxplots were generated using R version 3.2.2 (R Foundation for Statistical Computing; R studio 0.99.467) with ggplot2 package (Wickham et al., 2007, 2015).

### Total internal reflection fluorescence microscopy

Experiments in which septin filaments were polymerized in solution, reactions were incubated in plastic chambers were glued onto PEG-coated passivated coverslips for one hour to reach equilibria. Recombinant protein was diluted to 250 nM final concentration in reaction buffer (75 mM KCl, 50 mM Hepes pH 7.4, 1 mM BME, 0.1% methyl-cellulose, and 0.01% BSA).

For experiments on planar supported lipid bilayers, recombinant septins were incubated to a final concentration of 3 nM in buffer with 50 mM KCl, 50 mM Hepes pH 7.4, 1 mM BME, and 0.01% BSA. On lipid bilayers, septin polymerization was imaged immediately upon incubation.

Images were acquired utilizing a Nikon TiE TIRF system equipped with a solid-state laser system (15 mW, Nikon LUn-4), a Nikon Ti-82 stage, a 100x Plan Apo 1.49 NA oil lens, and Prime 95B CMOS camera (Photometrics). TIRF microscopy images were processed and analyzed using ImageJ (FIJI).

### Persistence length measurements

The cosine correlation of tangent angles along the filament lengths were assembled from single time points and fitted to an exponential to determine the persistence length using a MATLAB GUI previously published (Graham et al., 2014).

### Helical diagram generation

Amphipathic helical wheels were generated, and their net charge and hydrophobicity were determined using Heliquest (Gautier et al., 2008).

## Acknowledgements

We thank Danny Lew for generously providing *cdc12-6* and *shs1Δ* strains. This work was supported by NIH Grant R01GM130934 to A.S.G. B.L.W. was supported by the NIH Training Grant 2T32AI052080-16. K.S.C. was supported in part by a grant from the National Institute of General Medical Sciences under award T32 GM119999.

B.L.W. and K.S.C. designed and conducted the experiments. K.S.C. constructed new plasmids for protein expression. B.L.W. constructed strains for experiments. B.L.W. and A.S.G. wrote the paper with input from K.S.C.

---

**Movie S1. Wild-type and cdc12-6 septin filaments in solution**

TIRF movies of wild-type (left) and cdc12-6 (right) septin filaments formed in solution as in Figure 4D. Movies acquired and contrasted identically. Scale bar, 10 μm.

**Table S1.**
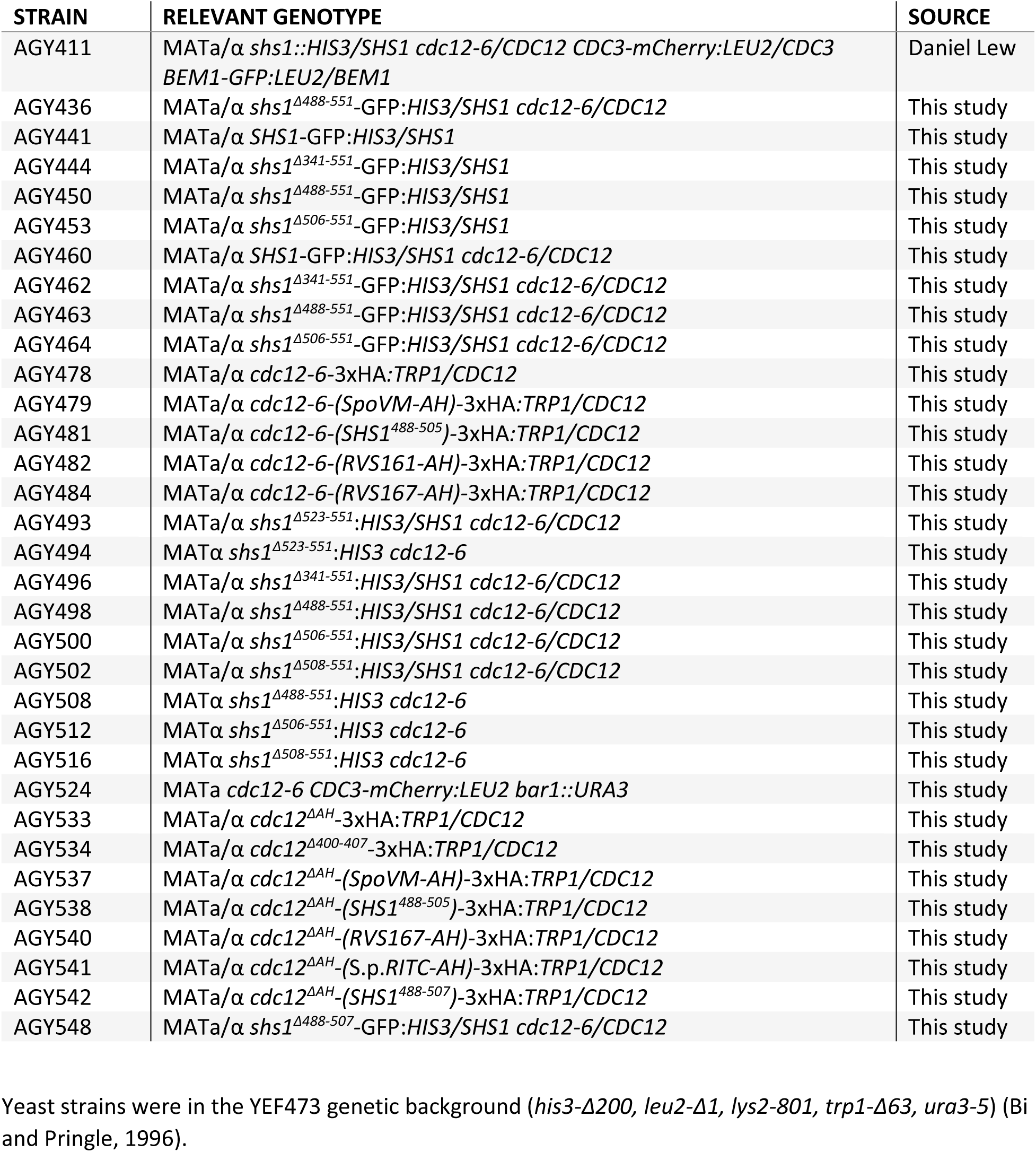
Yeast strains used in this study.

**Table S2.** Plasmids used in this study.

| NAME | DESCRIPTION | SELECTION MARKER |
| --- | --- | --- |
| AGB 548 | pMVB133 ScCDC3/ ScSHS1-GFP | CAM |
| AGB 710 | pMVB128 6-HIS-TEV-ScCDC12/ScCDC10 | AMP |
| AGB 1084 | pUC57 2xScShs1AH-GFP | AMP |
| AGB 1204 | pET-duet-6HIS-TEV-ScCDC12-6/ScCDC10 | AMP |

